# Supplementary motor area contributions to rhythm perception

**DOI:** 10.1101/2021.11.25.470060

**Authors:** Li-Ann Leow, Cricia Rinchon, Marina Emerick, Jessica A. Grahn

## Abstract

Timing is everything, but our understanding of the neural mechanisms of timing remains limited, particularly for timing of sequences. Temporal sequences can be represented relative to a recurrent beat (beat-based or relative timing), or as a series of absolute durations (non-beat-based or absolute timing). Neuroimaging work suggests involvement of the basal ganglia, supplementary motor area (SMA), the premotor cortices, and the cerebellum in both beat- and non-beat-based timing. Here we examined how beat-based timing and non-beat-based sequence timing were affected by modulating excitability of the supplementary motor area, the right cerebellum, and the bilateral dorsal premotor cortices, using transcranial direct current stimulation (tDCS). Participants were subjected to a sham stimulation session, followed an active stimulation session where anodal or cathodal 2mA tDCS was applied to the SMA, right premotor cortex, left premotor cortex, or the cerebellum. During both sessions, participants discriminated changes in rhythms which differentially engage beat-based or non-beat-based timing. Rhythm discrimination performance was improved by increasing SMA excitability, and impaired by decreasing SMA excitability. This polarity-dependent effect on rhythm discrimination was absent for cerebellar or premotor cortex stimulation, suggesting a crucial role of the SMA and/or its functionally connected networks in rhythmic timing mechanisms.

---

A fascinating human behaviour is the capacity to perceive the beat in sequences of temporal intervals (e.g., in music), even though beats are not necessarily indicated by distinguishing acoustic features. Beat perception, or the ability to sense a beat in rhythms, appears spontaneously in humans, without training (Winkler *et al.*, 2009). Beat perception is thought to engage relative timing mechanisms, in which the temporal intervals of a pattern are coded relative to each other (Essens & Povel, 1985; Yee *et al.*, 1994; Teki *et al.*, 2011a; Teki *et al.*, 2011b). This relative timing is often called ‘beat-based’ timing, because the intervals can be encoded relative to a regular, periodic beat interval. Beat-based timing improves accuracy during discrimination, synchronization, and reproduction of temporal sequences (Essens & Povel, 1985; Yee *et al.*, 1994; Patel *et al.*, 2005; Grahn & Brett, 2007; Chen *et al.*, 2008b). Relative timing stands in contrast to ‘absolute’ timing, also termed duration-based or non-beat-based timing, in which the absolute durations of temporal intervals are encoded individually in a stop-watch like manner (Teki *et al.*, 2011a).

An large body of functional neuroimaging studies have suggested involvement of the supplementary motor area (SMA), the basal ganglia, the premotor cortex, and the cerebellum in timing (e.g., Schubotz & von Cramon, 2001; Ullen *et al.*, 2003; Lewis *et al.*, 2004; Grahn & Brett, 2007; Chen *et al.*, 2008a; Bengtsson *et al.*, 2009; Teki *et al.*, 2011b). Determining the specific role of each area in different timing processes remains an active area of investigation. Neuroimaging studies find that the SMA and basal ganglia respond more to beat-based than non-beat-based than rhythms (Grahn & Brett, 2007; Grahn & Rowe, 2009; Teki *et al.*, 2011b; Geiser *et al.*, 2012; Grahn & Rowe, 2013; Li *et al.*, 2019). This pattern is consistent across tasks, including perceptual judgements (McAuley *et al.*, 2012), discrimination (Grahn & Brett, 2007), or attending to non-temporal aspects of the stimuli such as loudness (Geiser, Notter, & Gabrieli, 2012) and pitch (Grahn & Rowe, 2009).

Moreover, neuropsychological work in patients with Parkinson’s disease finds selective deficits in beat-based, but not non-beat-based timing (Grahn & Brett, 2009; Breska & Ivry, 2018). The premotor cortex and cerebellum appear to respond in both beat-based and non-beat-based contexts (Bengtsson *et al.*, 2005; Chen *et al.*, 2006), or respond more to non-beat-based than to beat-based contexts (Grahn & Rowe, 2009; Nozaradan et al., 2017; Teki et al., 2012). Moreover, patients with cerebellar degeneration show selective deficits in non-beat-based timing, despite intact beat-based timing (Grube *et al.*, 2010a; Breska & Ivry, 2018). It has thus been suggested that a functional network involving the SMA and basal ganglia subserves beat-based timing, whereas a functional network involving the cerebellum subserves absolute timing (e.g., Teki *et al.*, 2011b).

Importantly however, support for distinct neural processes subserving beat-based and non-beat based timing has mostly been supported by correlational neuroimaging evidence(Grahn & Brett, 2007; Grahn & Rowe, 2009; Teki *et al.*, 2011b; Geiser *et al.*, 2012; Grahn & Rowe, 2013; Li *et al.*, 2019), or by neuropsychological work in patient populations (Grahn & Brett, 2009; Grube *et al.*, 2010a; Breska & Ivry, 2018) who may have global deficits that influence task performance or compensatory changes over time. Relatively few studies employ perturbational methods in neurotypical humans, and such studies typically perturb only one or two brain areas implicated in beat perception (Malcolm *et al.*, 2008; Grube *et al.*, 2010b; Giovannelli *et al.*, 2014; Pollok *et al.*, 2017; Ross *et al.*, 2018b). Here, in neurotypical young adults, we examine how beat perception is affected by modulating multiple brain areas implicated in beat-based and non beat-based timing (i.e., supplementary motor area, left and right premotor cortex, and cerebellum), using transcranial direct current stimulation. Transcranial direct current stimulation (tDCS) is thought to modulate spontaneous neural firing and synaptic efficacy of neurons by altering the resting membrane potential, either increasing excitability through anodal stimulation, or decreasing excitability through cathodal stimulation (e.g., Bindman *et al.*, 1962; Lafon *et al.*, 2017). Therefore, tDCS has been proposed to have functionally specific effects by modulating activity of task-relevant neuronal networks (Bikson & Rahman, 2013). Given the large individual differences in beat perception ability (Grahn & McAuley, 2009; Grahn & Schuit, 2012; Sowiński & Dalla Bella, 2013), as well as large individual differences in tDCS responsivity (Chew *et al.*, 2015), we employed a within-subjects approach, where participants completed a placebo (sham) tDCS session followed by an active tDCS session whilst discriminating between rhythms which differentially engage beat-based timing. We hypothesized that a functional network involving the SMA supports beat-based timing: thus, increasing or decreasing SMA excitability was predicted to improve or impair discrimination of beat-inducing rhythms, and such effects should be more prominent for beat-inducing rhythms for non-beat-inducing rhythms. Moreover, as the cerebellum and PMC have been implicated (to varying degrees) in non-beat-based timing, then stimulation of these areas should have more prominent effects on non-beat-inducing rhythms than beat-inducing rhythms, although this prediction is supported to a greater degree for the cerebellum than for the premotor cortex.

## Method

### Participants

A total of 121 participants completed the experiment for course credit. Cerebellum anodal (n=14), cerebellum cathodal (n=16), left premotor anodal (n=15), left premotor cathodal (n=16), right premotor anodal (n=15) right premotor cathodal (n=15), SMA anodal (n=15), SMA cathodal (n=15). Participants were not pre-selected for music or dance training. Participants passed a safety screening for tDCS and gave informed consent. The Human Research Ethics Committee at Western University approved the study and all experiments were performed in accordance with relevant guidelines and regulations.

### Transcranial Direct Current Stimulation

Before behavioural testing, the scalp area overlying the stimulation site was located using the international electroencephalographic 10-20 system, as has been shown to be sufficient for tDCS using large electrodes as used here (Fregni *et al.*, 2006). Electrode montages were as follows. **SMA**: active electrode positioned 2 cm anterior to Cz, reference electrode positioned on contralateral orbit (Vollmann *et al.*, 2013). **Cerebellum**: active electrode positioned 3 cm right of the inion, reference electrode positioned on the right buccinator muscle (Galea *et al.*, 2009). **PMC**: active electrode positioned 2 cm anterior of C3 for left PMC, and 2cm anterior of C4 for right PMC (Nitsche *et al.*, 2003; Boros *et al.*, 2008), as neuroimaging studies suggest that the dorsal premotor cortex is located about 15–25 mm anterior to the primary motor cortex (C3, C4)(Picard & Strick, 2001). The reference electrode was positioned on contralateral orbit for both right and left PMC. Electrodes were secured using Velcro straps. For the active tDCS conditions, the current was gradually ramped up to the 2 mA level over 30 s upon commencing the rhythm discrimination task. The stimulation remained on during the task for a maximum of 40 minutes and was ramped down at the end of the stimulation period. For the sham tDCS conditions, the stimulation was similarly ramped up to 2 mA, but then ramped off over 30 s immediately after reaching 2 mA. This method achieves reasonable blinding in stimulation-naïve participants (Ambrus *et al.*, 2012). Stimulation was generated with a Dupel Stimulator (Dupel Ionophoresis System, MN) using two 4 x 6 cm rubber electrodes placed in saline-soaked sponges (current density of 0.04 mA/cm2; 0.9% NaCl) and highly conductive electrode gel (e.g., Signa Gel 40,000 micromhos/cm).

We employed a within-subjects approach to counter the large individual differences in beat perception ability (Grahn & McAuley, 2009; Grahn & Schuit, 2012; Sowiński & Dalla Bella, 2013), as well as large individual differences in tDCS responsivity (Chew *et al.*, 2015).

Each participant first completed a first sham tDCS condition in a first session, followed by active tDCS in the second session. In each session, during stimulation, participants completed the rhythm discrimination task (described below), with different rhythms used for each session to reduce practice effects. Participants completed both sessions on one visit, allowing us to eliminate participant drop-outs between visits. Time constraints made it unfeasible to have the active tDCS condition before the sham tDCS condition within a single-visit design, as effects can persist up to an hour post-stimulation (Nitsche & Paulus, 2000): waiting for tDCS effects to washout would require the visit to last more than 3 hours.

### Auditory Stimuli

We used rhythms known to differentially induce beat perception (Grahn & Brett, 2007; 2009; Grahn, 2012). These rhythms were created using integer ratio-related set of intervals (1:2:3:4). The shortest interval (i.e., 1) ranged from 225, 250, to 275 ms, selected on each trial and consistent for all rhythms in that trial. Sine tones (rise/fall times of 8 ms) sounded for the duration of each interval, ending 40 ms before the specified interval length to create a silent gap that demarcated the intervals. The other intervals in the rhythm were multiples of the shortest interval. For **beat-inducing rhythms**, the intervals were arranged in groups of four units (e.g., in the sequence 211314, an interval onset consistently occurs every four units). The rhythms were constructed to induce a regular perceptual accent at the beginning of each group of four units (Povel & Okkerman, 1981). cueing participants toe the regular beat structure, in which the beats coincided with the onset of each group (Essens, 1995). For **non-beat-inducing rhythms**, the intervals were not reliably grouped (e.g., 341211), and had no regular accent occurrence, thus not beat was induced.

**Figure 2.**
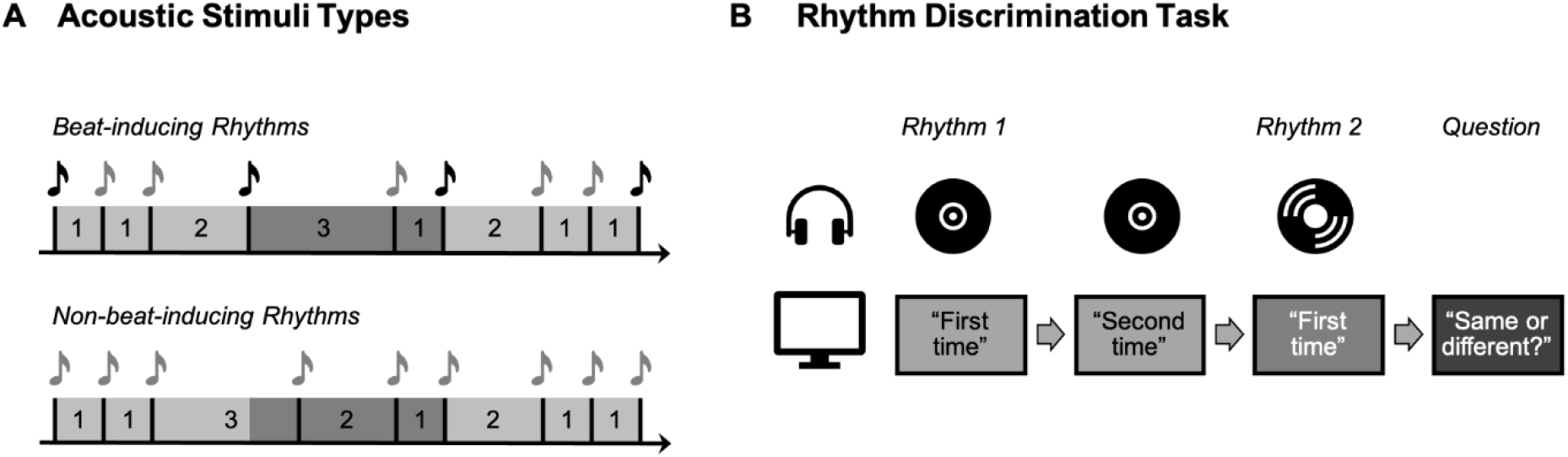
(A) Acoustic stimulus types: schematic example of the two types of rhythmic sequences used. Numbers denote the relative length of intervals in each sequence. 1 = 220-270 ms (selected on a per trial basis). Beat-inducing rhythms had intervals that could be grouped regularly (i.e., 1, 1, 2 = a group of 4, as are 3,1 and 2,1,1), and induced a percept of a beat. Non-beat-inducing rhythms could not be grouped regularly, and are more difficult to perceive a beat in. **(B) Rhythm Discrimination Task**: participants heard a rhythm twice, and then a second rhythm. Participants judged whether the third rhythm was the same as or different from the first rhythm.

### Task Procedure

The task was to judge whether the third presentation of a rhythm was the same as or different from the first two presentations. Rhythms were presented binaurally over Sennheiser headphones. The words ‘First time’, ‘Second time’, and ‘Same or different?’ were displayed on the screen during the first, second, and third rhythm presentations, respectively. On half of the trials (30 trials, 15 beat-inducing, 15 non-beat-inducing), the second rhythm played on the third presentation was the same as the first rhythm played on the first two presentations (non-deviant trials). On the other half of the trials (30 trials, 15 beat-inducing, 15 non-beat-inducing), the second rhythm played on the third presentation deviated from the first rhythm played on the first two presentations (deviant trials). The deviant sequences contained a transposition of intervals in the sequence (see Table 1). For example, the sequence 211413 can have as a possible deviant sequence 211431, in which the 3 interval and the 1 interval have been transposed. Only deviant sequences that were in the same category as the standard sequences were allowed. That is, a beat-inducing standard sequence could not have a non-beat-inducing deviant sequence, and a non-beat-inducing standard sequence could not have a beat-inducing deviant sequence. Participants indicated their response by pressing one key for ‘same’ and another key for ‘different’ on a computer keyboard. Between trials, there was a 2s interval of silence.

**Table 1.** Stimuli for non-deviant trials and deviant trials.

| <u>Non-deviant trials (first rhythm = second rhythm)</u> |  |  |  | <u>Deviant trials (first rhythm ≠ second rhythm)</u> |  |  |  |
| --- | --- | --- | --- | --- | --- | --- | --- |
| <u>Beat-Inducing</u> |  | <u>Non-Beat-Inducing</u> |  | <u>Beat-Inducing</u> |  | <u>Non-Beat-Inducing</u> |  |
| *First rhythm* | *Second rhythm* | *First rhythm* | *Second rhythm* | *First rhythm* | *Second rhythm* | *First rhythm* | *Second rhythm* |
| 22413 | 22413 | 11343 | 11343 | 41331 | 43131 | 11343 | 13143 |
| 31413 | 31413 | 23241 | 23241 | 43122 | 41322 | 23241 | 23214 |
| 31422 | 31422 | 33141 | 33141 | 211134 | 211314 | 33141 | 31341 |
| 43113 | 43113 | 41133 | 41133 | 211224 | 112224 | 41133 | 14133 |
| 112314 | 112314 | 214221 | 214221 | 211413 | 211431 | 214221 | 241221 |
| 112422 | 112422 | 321411 | 321411 | 223113 | 223131 | 321411 | 324111 |
| 221331 | 221331 | 323211 | 323211 | 311322 | 313122 | 323211 | 322311 |
| 222114 | 222114 | 421311 | 421311 | 312213 | 312231 | 421311 | 241311 |
| 411231 | 411231 | 1112412 | 1112412 | 1111431 | 1111413 | 1112412 | 1112142 |
| 422112 | 422112 | 1132212 | 1132212 | 1122114 | 1121124 | 1132212 | 1123212 |
| 2112231 | 2112231 | 1314111 | 1314111 | 1123113 | 1123131 | 1314111 | 1311411 |
| 2211114 | 2211114 | 2331111 | 2331111 | 1123122 | 1121322 | 2331111 | 2313111 |
| 3121113 | 3121113 | 3113121 | 3113121 | 2113113 | 2113131 | 3113121 | 3131121 |
| 3122112 | 3122112 | 3221112 | 3221112 | 4111131 | 4111113 | 3221112 | 2321112 |
| 3141111 | 3141111 | 4111221 | 4111221 | 4221111 | 4211211 | 4111221 | 1411221 |

The base interval of 1 lasted for either 225, 250, 275 msec. All other intervals in that sequence are multiplied by length chosen for the 1 interval.

Whilst undergoing sham stimulation, participants completed the first session of 60 trials which consisted of 30 trials with non-deviant rhythms (15 beat-inducing, 15 non-beat-inducing); and 30 trials with deviant rhythms (15 beat-inducing,15 non-beat-inducing). Trial order was randomised. The first session was immediately followed by the second second, where participants were subjected to active stimulation, and completed the second session of 60 trials (with similar task structure as described above). The same rhythms were used for the first and the second sessions, but the base interval length differed between sessions.

### Data Analysis

Task performance was quantified using and accuracy (percent correct) and sensitivity (*d’* scores)) (Stanislaw & Todorov, 1999) with larger *d’* scores indicating better discriminability.

### Statistical analysis

We chose to use Bayesian statistics to evaluate evidence for the alternative hypothesis and for the null hypothesis. Unlike p-values, Bayes factors do not tend to over-estimate the evidence against the null hypothesis (Gelman & Tuerlinckx, 2000; Wetzels *et al.*, 2011). Analyses were conducted in JASP (Version 0.13.1; JASP Team, 2020). The default Cauchy prior widths (0.707) values in JASP were used to quantify the relative evidence that the data came from the alternative versus a null model. Jeffreys’s evidence categories for interpretation were taken as the standard for evaluation of the reported Bayes Factors, in which the size of the Bayes factors are estimated as inconclusive(1–3), moderate (3–10), or strong (>10) evidence for the hypotheses tested.

To evaluate stimulation-induced changes in discrimination performance across different stimulation sites and polarities, Rhythm (non-beat-inducing, beat-inducing) x Stimulation (sham, stimulation) x Polarity (anodal, cathodal) x Site (SMA, cerebellum, right PMC, left PMC) Bayesian ANOVAs were run on *d’* and percent correct. Analyses estimated the evidence for including each effect across matched models via estimating inclusion Bayes factor (BF_incl_) for each effect. Analyses also estimated the evidence for excluding each effect by estimating an exclusion Bayes factor (BF_excl_) for each effect. Where applicable, simple effects analyses were used to follow up interactions.

## Results

### Effect of tDCS on rhythm discrimination

#### Rhythm discrimination performance

Replicating previous results (Grahn & Brett, 2009; Grahn, 2012), *d’* and percent correct scores were higher for beat-inducing than for non-beat-inducing rhythms, indicating better discrimination for beat inducing rhythms [*d’*: beat-inducing: 1.76±0.08, non-beat-inducing: 1.16±0.06; percent correct: beat-inducing= 76.6 ± 1.1%, non-beat-inducing= 69.1±0.9%; main effect of rhythm type BF_incl_ = 6.27e+28]. Figures 3 and 4 respectively show *d’* and percent correct scores from the sham condition and stimulation conditions. Generally, patterns in percent correct scores appear similar to *d’* scores, as in previous work (Grahn, 2012).

**Figure 3.**
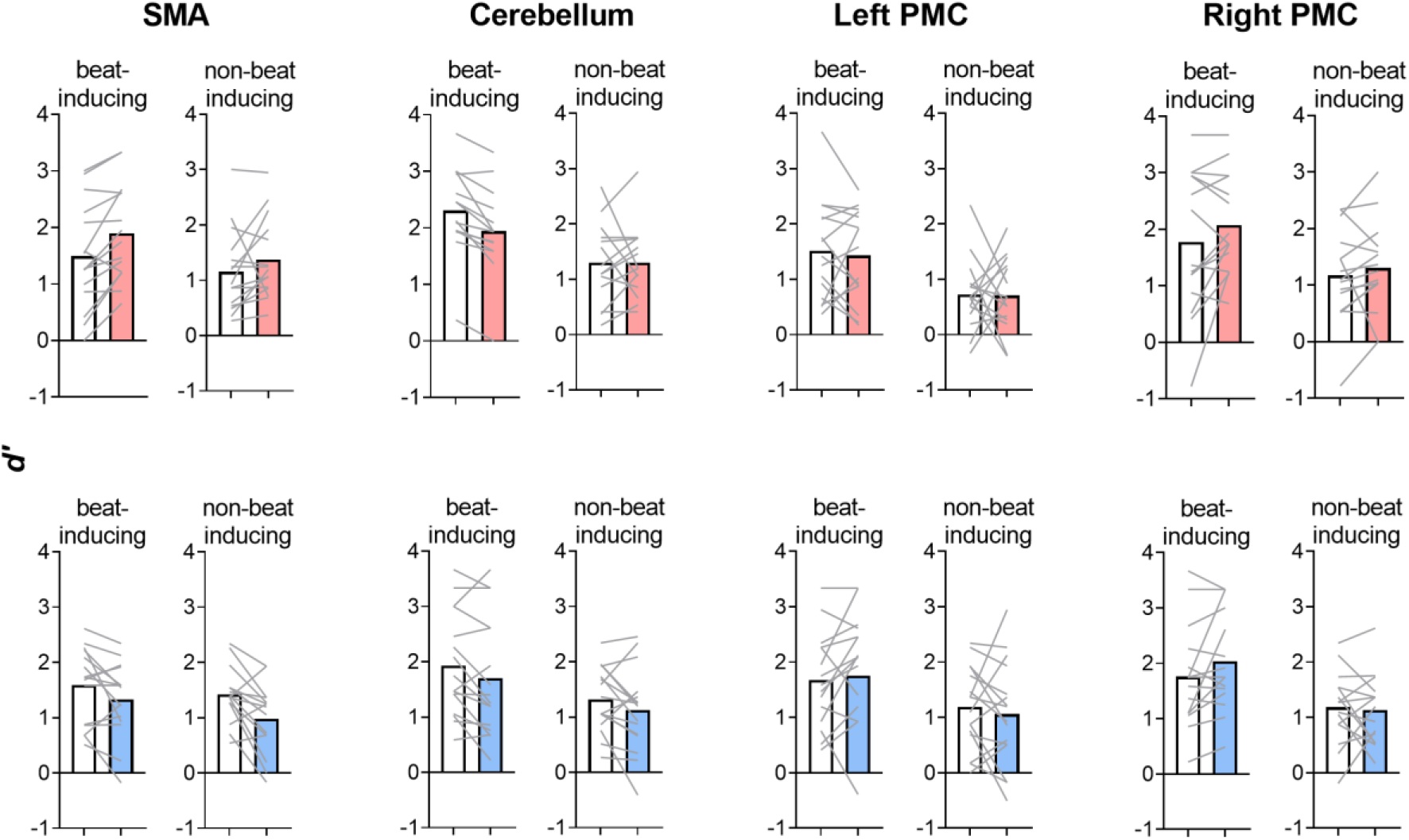
*d’* from the sham stimulation session (white bars) and the active stimulation session (coloured bars: pink = anodal tDCS, blue = cathodal tDCS), with beat-inducing and non-beat-inducing rhythms. There is moderate evidence for polarity-dependent effects in the SMA, inconclusive evidence for polarity-independent effects in the cerebellum, and moderate evidence *against* any effects of stimulation on the left and right PMC.

**Figure 4.**
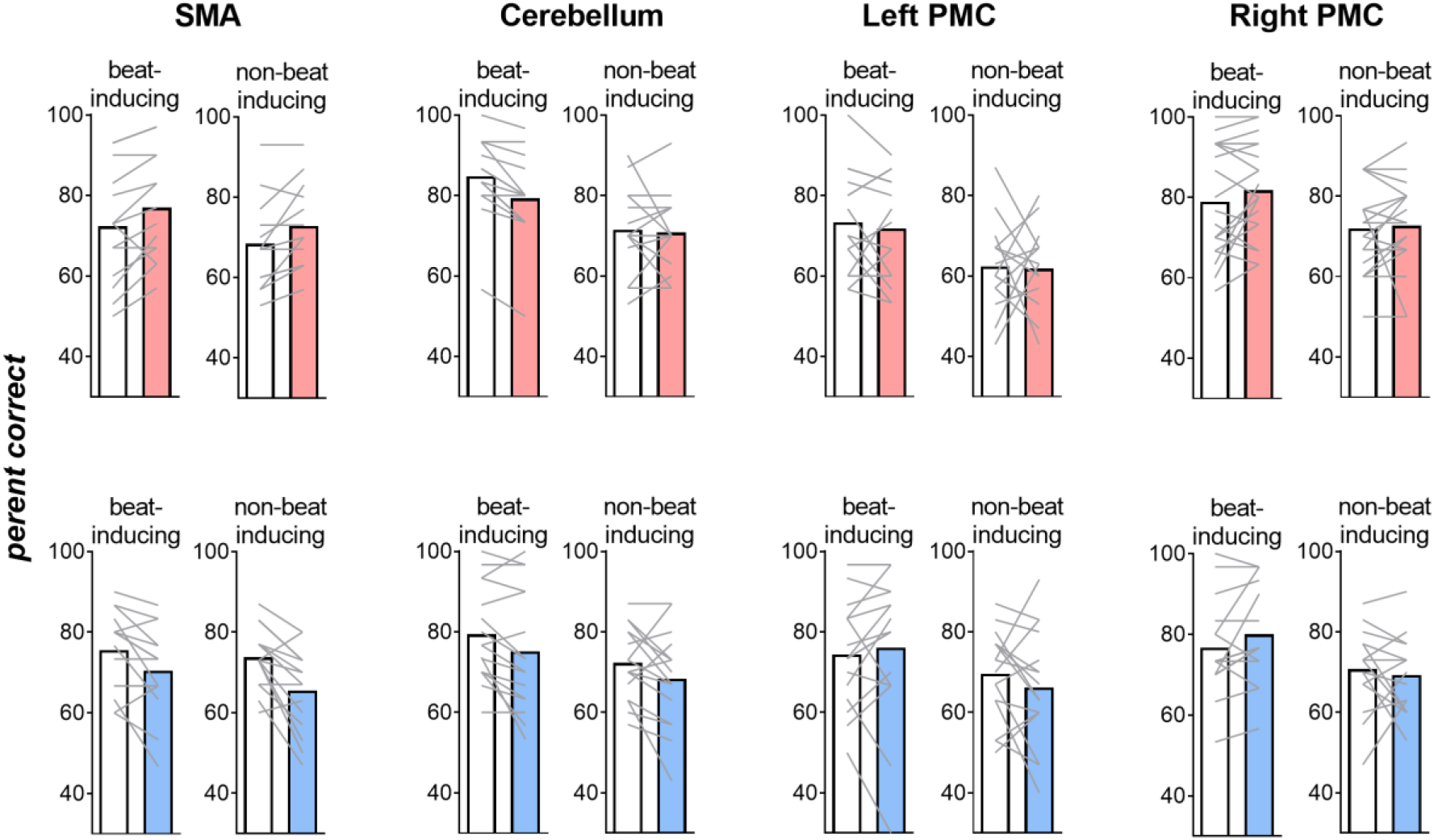
Response accuracy from the sham stimulation session (white bars) to the active stimulation session (coloured bars, where pink indicates anodal tDCS whereas blue indicates cathodal tDCS), as indicated by percent of trials correct for anodal (top panels) and cathodal (bottom panels) tDCS conditions, with beat-inducing and non-beat-inducing rhythms.

Stimulation affected rhythm discrimination performance differently depending on the stimulated brain area and stimulation polarity, as shown by moderate Stimulation x Site x Polarity interactions [*d’*: BF_incl_ = 3.019, percent correct: BF_incl_ = 2.574]. Follow-up Rhythm (non-beat-inducing, beat-inducing) x Stimulation (Sham, Stimulation) x Polarity (Anodal, Cathodal) Bayesian ANOVAs were therefore run separately for each stimulation site.

For SMA, there were strong Stimulation (Sham, Anodal) x Polarity (Anodal, Cathodal) interactions [*d’*: BF_incl_ = 298.500, percent correct: BF_incl_ = 288.447]. Follow-up simple effects analyses (i.e., Rhythm x Stimulation Bayesian ANOVAs) run separately for the anodal and cathodal conditions showed that there was moderate evidence for anodal SMA stimulation improving discrimination performance compared to sham (main effect of stimulation for *d’*: BF_incl_ = 4.060, percent correct: BF_incl_ = 3.679 (see Figure 3 and 4), and strong evidence for cathodal SMA stimulation worsening discrimination performance compared to sham (main effect of stimulation for *d’*: BF_incl_ = 16.732, percent correct: BF _incl_ =17.243). Compared to sham, anodal SMA tDCS improved discrimination for metric simple rhythms (*d’*: sham = 1.49+/−0.24, anodal = 1.89+/−0.22; percent correct: sham = 72.6+/−3.57%, anodal = 77.8+/−3.39) as well as for metric complex rhythms (*d’*: sham = 1.15+/−0.19; anodal= 1.37+/−0.18, percent correct: sham = 69.33+/−2.91%, anodal = 72.13+/− 2.63%). Cathodal tDCS impaired discrimination for both metric simple rhythms (d’: sham = 1.59+/− 0.17, cathodal = 1.33+/−0.17: percent correct: sham = 75.5+/−2.5%, cathodal = 70.4+/−2.8%) and metric complex rhythms (d’: sham = 1.42+/−0.12, cathodal: 0.98+/−0.16); percent correct: sham = 73.7+/−1.9%, cathodal = 65.5+/−2.6%).

For cerebellum, stimulation tended to worsen discrimination performance [*d’*: Sham = 1.70+/−0.13, stimulation = 1.51+/−0.13, main effect of stimulation, BF_incl_ = 1.941; percent correct: Sham = 76.8+/−1.9%, Stim = 73.3+/−1.9, BF_incl_ = 9.102]. The effect of stimulation appeared to be polarity-independent (moderate evidence against the stimulation by polarity interaction [*d’*: BF_excl_ = 3.710, percent correct: BF_excl_ = 2.985].

For the left premotor cortex, there was little evidence for an effect of stimulation, as there was moderate evidence *against* including the main effect of stimulation [*d’*: BF_excl_ = 5.175; percent correct: BF_excl_ = 4.466] and the stimulation x polarity interaction [*d’*: BF_excl_ = 3.614, percent correct: BF_excl_ = 3.913]. Similarly for the right premotor cortex condition, there was evidence against including the main effect of stimulation [*d’*: BF_excl_ = 2.315; percent correct: BF_excl_ = 7.347e −8] and the stimulation x polarity interaction [*d’*: BF_excl_ = 3.592, percent correct: BF_excl_ = 3.713].

## Discussion

We provide initial evidence suggesting that discrimination of beat-inducing and non-beat-inducing rhythms can be improved by increasing SMA excitability, and impaired by decreasing SMA excitability. This polarity-dependent effect on discrimination of beat-inducing rhythms was not evident for cerebellar or premotor cortex stimulation, suggesting a functional role of the SMA and/or its functionally connected networks in rhythm timing. However, contrary to our specific predictions, the effect of SMA stimulation was not unique to beat-inducing rhythms but instead altered performance similarly for both beat-inducing and non-beat-inducing rhythm. Interestingly, cerebellar stimulation reduced discrimination performance regardless of stimulation polarity, but not selectively for non-beat-inducing rhythms, in contrast to our predictions.

### How might the SMA support beat-perception?

Here, we found that modulating SMA excitability altered performance on the rhythm discrimination task in a polarity-dependent fashion. These findings support the idea that the SMA and the basal ganglia are involved in maintaining an internal representation of beat intervals (Grahn & Brett, 2007; Grahn & Rowe, 2009), facilitating performance on the rhythm discrimination task. Our results are broadly consistent with findings of greater SMA-basal ganglia activation during the processing of beat-inducing rhythms compared to non-beat-inducing rhythms (Grahn & Brett, 2007; Grahn & Rowe, 2009). It has been suggested that the SMA networks help to form forward temporal predictions (Macar *et al.*, 2004). The role of the SMA in beat-maintenance is consistent with evidence in synchronization-continuation tasks where participants are asked to synchronize movements to external stimuli and then continue synchronization upon withdrawal of the external stimuli: here, the SMA tends to be activated during the continuation phase, and not the synchronization phase (Rao *et al.*, 1997; Lewis *et al.*, 2004). Similarly, patients with SMA lesions also show a selective deficit in the continuation phase but not the synchronization phase of the synchronization-continuation task (Halsband *et al.*, 1993). Recent proposals have suggested that the SMA is tuned to anticipate an upcoming beat interval, and sends signals to the dorsal striatum, helping the striatum generate internally generated representations of the beat cycle, which in turn activates new SMA neural subpopulations via the thalamus (Cannon & Patel, 2021). Future studies might test this hypothesis via recordings in SMA and basal ganglia, perhaps in patients with electrocorticography and deep brain stimulator implants.

### Effect of premotor cortex tDCS on rhythm discrimination performance

The premotor cortex is thought to be engaged in planning, selection, and control motor programs based on external events (Picard & Strick, 2001). The finding that stimulation of the left or right premotor cortex did not interfere with rhythm discrimination performance is consistent with the idea that the premotor cortex plays a primary role in synchronization and control of motor programs in response to external events, rather than beat perception per se. Although some findings implicate a role of the premotor cortex in beat-based timing (Chen *et al.*, 2006; Chen *et al.*, 2008a; b; 2009), all of these studies involve the synchronization of rhythms using repetitive TMS. Synchronization requires participants to synchronize movements to the onset of each tone of a rhythm, or to each beat in the rhythm. This not only requires participants to encode and maintain the beat interval, but also to produce a synchronized motor response, evaluate the accuracy of that response after each tap, and correct the timing of inaccurate taps. Indeed, the greater difficulty of synchronizing to non-beat rhythms (and consequently greater demands on motor planning, evaluation, and error correction) might have resulted in the increased premotor cortex activation when synchronizing to non-beat-rhythms than beat rhythms (Chen *et al.*, 2008b). The notion that the premotor cortex plays a general role in synchronization of movements to external stimuli is supported by an increasing body of evidence showing that non-invasive stimulation of the premotor cortex modulates synchronization performance, when synchronizing to isochronous cues (Doumas *et al.*, 2005; Del Olmo *et al.*, 2007; Malcolm *et al.*, 2008; Pollok *et al.*, 2008; Bijsterbosch *et al.*, 2011; Ruspantini *et al.*, 2011), when adjusting for changes in cue onsets (Bijsterbosch *et al.*, 2011; Kornysheva & Schubotz, 2011; Ruspantini *et al.*, 2011), or when tapping to rhythms which differentially engage beat perception (Kornysheva & Schubotz, 2011; Giovannelli *et al.*, 2014). These findings show distinct effects of stimulation on aspects of synchronization (e.g., tempo-matching versus phase-matching), and different effects on dorsal versus ventral premotor cortex stimulation, or left versus right premotor cortex stimulation. A detailed discussion of these findings is outside the scope of this study. To the best of our knowledge, there are no papers published in peer-reviewed journals demonstrating effects of stimulating the premotor cortex on beat perception tasks that do not require motor synchronization to external stimuli in humans. One study currently published on a preprint server has examined how stimulation of premotor cortex affects capacity to perceive changes in tempo in music (Ross *et al.*, 2018a). However, music contains many redundant cues that signal beat onsets which aid the perception of tempo changes: effects of premotor cortex stimulation in this study might thus not be directly related to beat perception in temporal intervals without redundant cues. To elucidate the role of the premotor cortex in perceiving and synchronizing to the beat, future studies should examine effects of modulating premotor cortex excitability on the processing of beat and non-beat-based rhythms using both perceptual tasks and motor synchronization within the same subjects.

### Effect of cerebellar tDCS on rhythm discrimination performance

Cerebellar tDCS resulted in lower discrimination accuracy (i.e., percent correct scores) regardless of stimulation polarity, with both beat and non-beat rhythms, although this effect appeared more robust for percent accuracy than discrimination sensitivity (*d’)*. The polarity-independent effect on discrimination accuracy is perhaps unsurprising given the increasing numbers of studies that show polarity-independent effects of anodal and cathodal cerebellar tDCS on working memory (Ferrucci et al., 2008; Van Wessel et al., 2016), motor control and learning (Shah et al., 2013; Verhage et al., 2017), motor memory retention (Taubert et al., 2016), and conditioned eyeblink responses (Beyer et al., 2017). Indeed, one recent meta-analysis found no evidence for a polarity-dependent effect of cerebellar tDCS (Oldrati & Schutter, 2018). Polarity-dependent effects of tDCS on cortex do not necessarily generalize to the cerebellum, as the organization of cerebellar neurons differs fundamentally from that of the cortex (van Dun et al., 2016; Woods et al., 2016).

A few interpretations are possible for how cerebellar tDCS impaired discrimination accuracy here. First, cerebellar tDCS might have induced impairments in working memory, similar to previous findings (Van Wessel et al., 2016)(Ferrucci et al., 2008). In the rhythm discrimination task used here, working memory is required to remember and compare the first rhythms with the test rhythms. Second, cerebellar tDCS might have impaired absolute timing processes, consistent with previous findings of worsened absolute timing (Grube *et al.*, 2010b). Both interpretations may be true.

Judicious experimental designs which explicitly manipulate working memory load and/or stimuli which differentially engage absolute timing might provide evidence for or against these two possible interpretations.

### Limitations

An important limitation in the current work is the lack of counterbalancing of the order of sham and active tDCS: the sham tDCS session always preceded the active tDCS session. However, previous work using these rhythms has shown no reliable improvement between sessions (the familiarization trials are sufficient to induce steady-state performance (Cameron *et al.*, 2016)), thus practice effects appear minimal for this task. Moreover, only the group receiving anodal SMA tDCS improved rhythm discrimination performance from the first sham tDCS session to the second active tDCS session. No improvement in discrimination performance was evident in any other group, and cathodal SMA tDCS impaired rhythm discrimination. This polarity-dependent effect suggests order effects cannot be the explanation. A general learning effect would manifest for all other sites (left PMC, right PMC, cerebellum). Finally, previous studies which test the effect of SMA stimulation do not show effects of SMA tDCS on learning (Foerster *et al.*, 2013). It thus seems likely that SMA tDCS modulated capacity to discriminate rhythms, rather than *learning* to discriminate rhythms.

A second limitation is the difficulty in blinding tDCS administration. Blinding in tDCS can be challenging even using double-blind designs where one experimenter conducts electrode placement and initiates stimulation and a different experimenter administers the behavioural task (O’connell *et al.*, 2012). Here we used a single-blind design. Although our protocols of employing sham tDCS first followed by active tDCS could have unblinded participants (Turi *et al.*, 2019), Participants likely remained blinded to stimulation *polarity*, therefore their expectations are unlikely to have resulted in polarity-dependent effect of stimulation the SMA. We also did not predict the effect of cerebellar tDCS impairing rhythm discrimination performance regardless of stimulation polarity: this effect also seems unlikely to have resulted from participant expectations. Finally, the selective polarity-dependent effect on SMA but not for right and nor left premotor cortex conditions provides robust evidence for a genuine effect of SMA stimulation on rhythm discrimination performance, as any unblinding effects should apply equally across these nearby brain regions. Therefore, (for a review, see Ridding & Ziemann, 2010; Chew *et al.*, 2015; López-Alonso *et al.*, 2015; e.g.,Filmer *et al.*, 2020)this work presents evidence suggesting that SMA tDCS modulates rhythm discrimination performance.

### Summary

Neuroimaging and neuropsychological evidence implicate a role of a functional network encompassing the supplementary motor area and the putamen in beat perception. Here, we show that non-invasive stimulation of the supplementary motor area can have polarity-dependent effects on discrimination of auditory rhythm. Although the current evidence implicates a role of the supplementary motor area in beat perception, exactly how the SMA interacts with other brain areas during beat perception remains unclear. Exploring this question by combining perturbational methods such as brain stimulation with methods that afford both high temporal and spatial resolution (e.g., magnetoencephalography) could elucidate the mechanisms through which regions implicated in processing time interact and contribute to beat perception.

